# A closed-loop cell therapy engineered to autonomously secrete Activin A inhibitor protects from fibrodysplasia ossificans progressiva

**DOI:** 10.64898/2026.02.11.705403

**Authors:** Pratik Koirala, Ziyu Chen, Mengfan Wu, David Maridas, Ashley Siegel, Chang Liu, Samerender Hanumantharao, Yuji Mishina, Vicki Rosen, Shailesh Agarwal

## Abstract

Engineered cell therapies present an opportunity for endogenous, site-specific production of therapeutic agents. Here we describe a closed-loop cell therapy which secretes an inhibitor of Activin A, ActR2A-Fc, upon exposure to Activin A. We demonstrate in vivo therapeutic efficacy of this approach in a mouse model of fibrodysplasia ossificans progressiva (FOP), a morbid condition in which patients develop extensive heterotopic bony lesions in response to aberrant sensitivity to Activin A through a mutation in the type I BMP receptor ACVR1 (ACVR1 R206H). To blunt Activin A activity, we designed a transposon plasmid containing the transgene encoding ActR2A-Fc, with expression controlled by the BMP-responsive element (BRE). In cells containing the causative mutation, the BRE is pathologically activated upon exposure to Activin A. FOP-derived marrow cells modified with the BRE-ActR2AFc plasmid exhibited the desired closed-loop functionality, with increased ActR2A-Fc expression upon exposure to Activin A and reduced expression upon withdrawal of Activin A. Engineered marrow cells secreted bioactive ActR2A-Fc, and bone marrow transplantation of FOP marrow cells engineered with the BRE-ActR2AFc transposon into same-sex FOP mice resulted in absence of heterotopic bony lesions. Experiments with labeled, engineered FOP marrow cells verified trafficking of the therapeutic cells to sites at risk for FOP. These data provide proof-of-concept for the therapeutic utility of engineered cell therapy for the treatment of FOP.

**Significance statement:** In this study, we describe our development of an autologous, closed-loop cell therapy which can migrate to sites of tissue injury and locally secreting an inhibitor of Activin A. Through our use of an Activin A-responsive promoter to drive expression of the recombinant Activin A inhibitor, this engineered cell therapy exhibits closed-loop behavior and effectively prevents heterotopic bone formation in a mouse model of fibrodysplasia ossificans progressiva (FOP). We believe that the findings in this manuscript impactful beyond FOP, and provide a blueprint for the development of marrow-derived cell therapies across the disease spectrum.

## Introduction

Drug delivery systems that enable local, spatially restricted delivery of therapeutic agents to the affected disease site offer the possibility of administering larger doses than would be tolerated systemically. Cell therapies offer an opportunity for spatiotemporally restricted drug delivery by taking advantage of three unique characteristics. First, cell therapies synthesize the pharmacologic agent, thereby obviating exogenous drug delivery. Second, the cell utilized are able to traffic to involved sites of tissue injury through the local presence of recruitment cytokines such as stromal-derived factor 1 (SDF1)^1,2^; as a result, their function can be spatially-restricted. Finally, cell therapies are activated only when engaged with the offending disease process. This feature can enable cell therapies to activate in a pseudotemporal fashion^3^, in which time is defined by the presence of the disease.

Chimeric antigen receptor T-cell therapy (CAR T-cell therapy) presents one of the most well-recognized strategies for spatiotemporally restricted treatment^4^. Outside of treatment for malignancy, cell therapy adoption specifically for drug delivery has been unrealized. There are myriad conditions, many of them driven by local inflammation, which would benefit from cell therapies able to traffic to sites of tissue injury, activate locally, and secrete a recombinant therapy. Here we report on the development of a cell therapy which is suited to treat fibrodysplasia ossificans progressiva (FOP)^5^, a rare condition in which patients develop painful heterotopic bony lesions at sites of tissue injury. Patients with FOP harbor a germ-line mutation in the bone morphogenetic protein (BMP) receptor *ACVR1*; this mutation (*ACVR1 R206H*) renders fibro/adipogenic progenitors (FAP) pathologically susceptible to osteogenic differentiation^6,7^ upon exposure to Activin A^8^, a member of the TGFβ superfamily. Activin A is a major component of the inflammatory response. The concentration of circulating activin A has been shown to increase during inflammation^9.^

Recent therapeutic strategies for FOP have demonstrated notable progress though the two leading approaches. Treatment with an anti-Activin A antibody, garetosmab, has shown high efficacy and favorable tolerability in clinical trials, leading to strong patient adherence in late-stage studies^10^. In contrast, the approved oral therapy palovarotene (Sohonos)^11^, which acts on downstream signaling, has seen limited clinical adoption due to concerns over its modest efficacy and significant side-effect profile. This highlights the ongoing need for FOP therapies that combine efficacy with long-term safety and patient tolerability.

Using transposon technology^12^, we have engineered a closed-loop cell therapy which is indefinitely capable of secreting an Activin A inhibitor upon exposure to Activin A, using an Activin A-inducible promoter. This cell therapy, designed to be an autologous marrow replacement, traffics to sites of inflammation and expresses the Activin A inhibitor, ActR2A-Fc, using the bone morphogenetic protein responsive element (BRE) promoter^8^ which is uniquely activated by Activin A in cells harboring the causative *ACVR1 R206H* mutation. Despite high levels of marrow reconstitution with our cell therapy, we show absence of systemically detectible drug levels. Marrow cells are recruited to sites of injury and there is near complete absence of heterotopic bone in FOP mice which receive the engineered cell therapy. While systemic drug delivery has demonstrated therapeutic promise for FOP, the requirement for chronic administration raises potential concerns for off-target effects and long-term toxicity. Our findings suggest that spatiotemporal drug delivery via genetically modified cell therapies provides a compelling alternative strategy. This approach, which restricts therapeutic protein expression to sites of injury and avoids systemic detection, may offer an improved safety profile for managing localized disease flares.

## Results

### Design of closed-loop plasmid targeting Activin A

We designed a transgenic plasmid which would activate gene expression in cells upon activation of BMP signaling using the BMP-responsive element (BRE) promoter. The plasmid includes 2 BRE repeats. To specifically target Activin A as the pathologic ligand responsible for BMP signaling in FOP cells carrying the *ACVR1 R206H* mutation, our plasmid included the transgene encoding the Activin-binding receptor, ActR2A-Fc (Fig 1A). This protein consists of the signal peptide and functional domain of the human ActR2A protein from Met1-Pro135 followed by the IEGRMD sequence and Fc tag from human IgG1. The closed-loop regulatory system for ActR2A-Fc expression responds to fluctuating Activin A levels. During inflammation, elevated Activin A concentration activates the BMP-responsive element (BRE) promoter, driving the expression of ActR2A-Fc. The secreted ActR2A-Fc sequesters extracellular Activin A, lowering its bioavailability and attenuating further ActR2A-Fc expression, thus creating a feedback loop. Transfected bone marrow cells expressing ActR2A-Fc in response to Activin A serve as a therapeutic intervention for fibrodysplasia ossificans progressiva (FOP) caused by the *ACVR1 R206H* mutation. By trapping Activin A, the ActR2A-Fc produced by these therapeutic cells inhibits downstream Smad1/5 pathway activation, effectively suppressing pathological osteogenesis and reducing heterotopic bone formation (Fig 1B).

**Figure 1.**
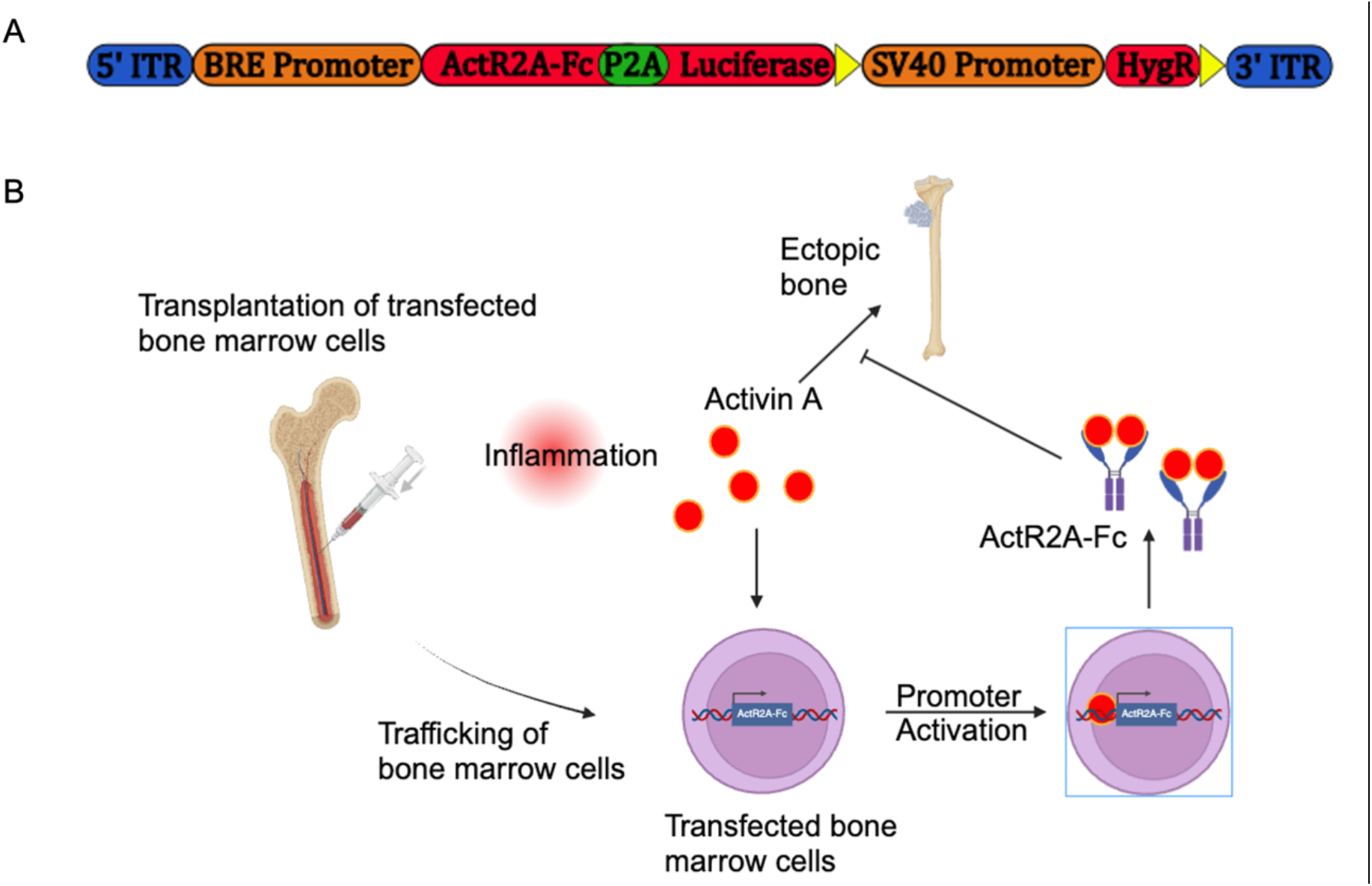
Closed-loop strategy targeting Activin A. (A) Schematic representation of the BRE-ActR2A-Fc construct. (B) Proposed model for transfected bone marrow cells-based therapy in fibrodysplasia ossificans progressiva (FOP) mice.

### Validation of expression and secretion of ActR2A-Fc

HEK293 cells transfected with the plasmid exhibited BMP2-dependent ActR2A-Fc expression (Fig 2A, B). Similarly, transfected HEK293 cells also exhibited BMP2-dependent ActR2A-Fc secretion based on ELISA (Fig 2C). Upon validating the BMP-responsiveness of the plasmid in HEK293 cells, we proceeded with transfection into wildtype (WT) mouse bone marrow cells. In similar fashion, WT bone marrow cells transfected with the plasmid exhibited BMP2-dependent ActR2A-Fc expression (Fig 2D). Similarly, transfected bone marrow cells also exhibited BMP2-dependent ActR2A-Fc secretion based on ELISA (Fig 2E).

**Figure 2.**
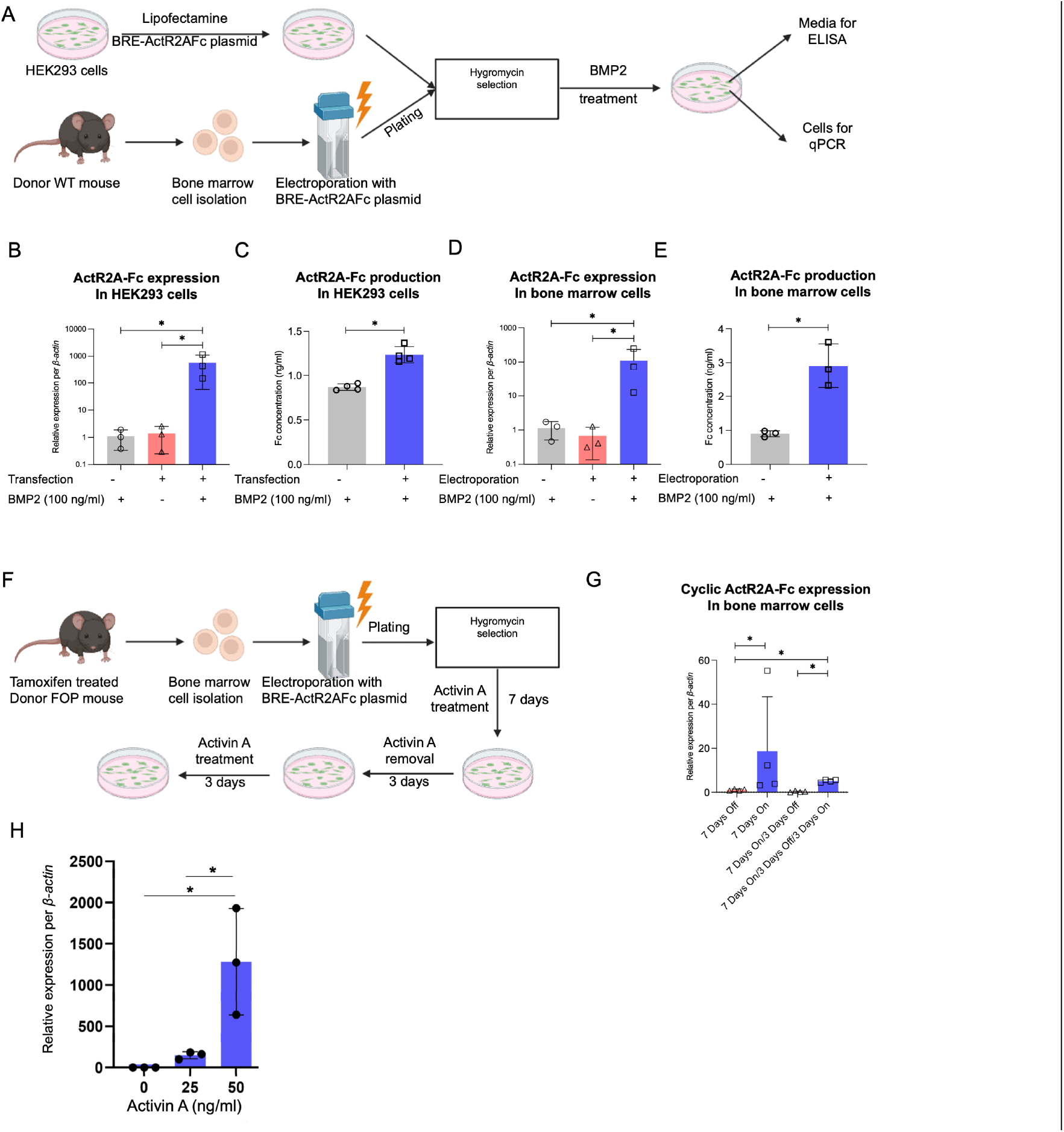
Expression and secretion of ActR2A-Fc. (A) Schematic of the in vitro ActR2A-Fc expression and production assay in HEK293 and WT bone marrow cells. (B) qPCR results showing BMP2-responsive ActR2A-Fc expression in transfected HEK293 cells. (C) ELISA results showing BMP2-responsive ActR2A-Fc production in transfected HEK293 cells. (D) qPCR results showing BMP2-responsive ActR2A-Fc expression in transfected WT mouse bone marrow cells. (E) ELISA results showing BMP2-responsive ActR2A-Fc production in transfected WT mouse bone marrow cells. (F) Schematic for the in vitro assay to verify the reversibility of ActR2A-Fc expression in transfected FOP mouse bone marrow cells. (G) Transfected FOP mouse bone marrow cells displayed significantly elevated ActR2A-Fc expression after 7 days of Activin A treatment. Expression returned to basal levels upon removal of Activin A and increased again after re-addition of Activin A. (H) Transfected FOP bone marrow cells displayed dose-dependent activation of ActR2A-Fc with increasing concentration of Activin A. Each dot in graphs represents a single biological replicate. Statistical significance was established by performing Welsh’s t-test between the means of the groups. * is p<0.05.

Next, we sought to validate the closed-loop functionality of our plasmid. To do this, we directed our efforts to FOP mouse bone marrow cells as these were the desired target cells for our plasmid in vivo. We designed an in vitro experiment to simulate the exposure, withdrawal, and re-exposure to the Activin A ligand (Fig 2F). We found that upon exposure to Activin A, transfected bone marrow cells exhibited the desired upregulation of ActR2A-Fc expression. This was attenuated upon withdrawal of Activin A, but then re-activated upon re-exposure to Activin A (Fig 2G). The reactivation was not as robust as the first activation; however, the second activation was after 3 days in the media compared to 7 days for the first activation, which might have led to lower expression. The expression of ActR2A-Fc in FOP bone marrow derived cells was Activin A dose-dependent. No activity was seen in the absence of Activin A. The expression significantly increased with 50 ng/ml Activin A (Fig 2H).

### Validation of bioactive ACVR2A-Fc production

Upon validation of our genetic construct, we verified the bioactivity of secreted ActR2A-Fc recombinant peptide using bone marrow cells from the FOP mice. Bone marrow cells from FOP mice were divided into two groups: transfected and untransfected. Both groups were treated with Activin A, and conditioned media were collected after 7 days (Fig 3A – schematic). We observed an increase in osteogenic gene expression, including Runx2, Bglap, Sox9 and Col10a1 when FOP mouse-derived bone marrow cells were treated with Activin A and grown in presence of conditioned media from untransfected bone marrow cells (Fig 3B-E). In contrast, osteogenic gene expression was notably reduced when FOP mouse-derived bone marrow cells treated with Activin A were grown in presence of conditioned media from transfected bone marrow cells (Fig 3B-E). To test whether ActR2A-Fc may reduce the effect of other ligands known to bind ActR2A, we performed experiments with BMP9, a ligand which binds ActR2A. A similar trend was seen when WT bone marrow cells were treated with BMP9 and grown in presence of ActR2A-Fc containing conditioned media (Fig 3F-I). This further demonstrates that ActR2A-Fc may bind to ligands which are already known to bind ActR2A and may present local off-target ligand inhibition.

**Figure 3.**
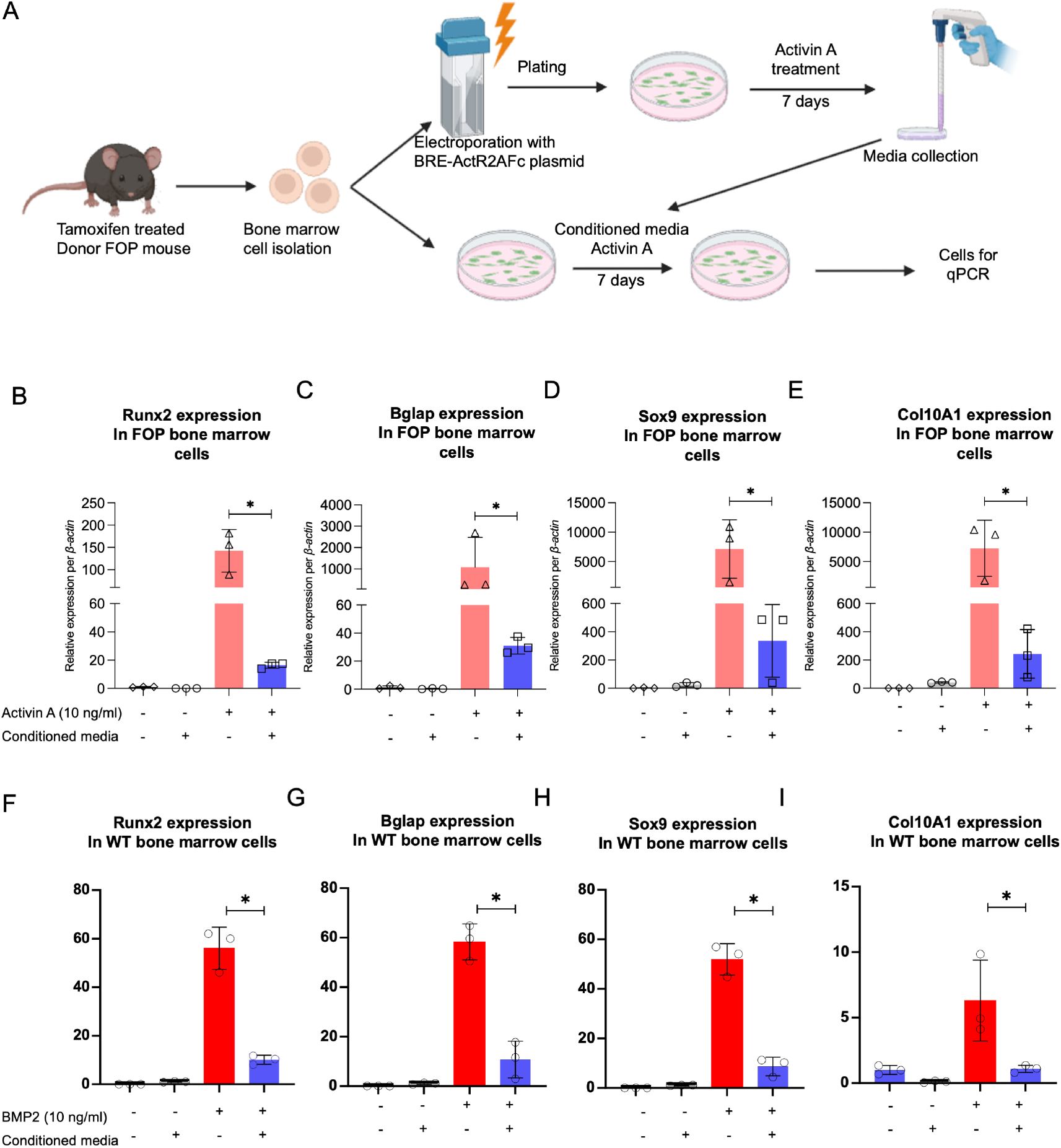
In vitro bioactivity of ActR2A-Fc secreted from FOP mouse derived bone marrow cells. (A) Schematic for in vitro validation of ActR2A-Fc bioactivity. (B-E) qPCR results showing expression of osteogenic genes (Runx2, Bglap, Sox9, Col10A1) in FOP mouse derived bone marrow cells. (F-I) qPCR results showing expression of osteogenic genes (Runx2, Bglap, Sox9, Col10A1) in WT mouse derived bone marrow cells. Each dot in graphs represents a single biological replicate. Statistical significance was established by performing Welsh’s t-test between the means of the groups. * is p<0.05.

### Transplantation of transfected bone marrow into FOP mice

Next, we proceeded with in vivo validation of our construct’s therapeutic efficacy. We followed a one-step marrow transfection procedure; marrow was isolated from tamoxifen-induced mT/mG FOP mice and transfected using electroporation (200 V/5 msec/1 pulse) (Fig 4A). We verified that immediately after and 1 week after electroporation, the ratio of CD45+ (hematopoietic) and PDGFRα+ (mesenchymal cell) populations remained similar in the untransfected and transfected groups (Fig 4B). Following electroporation, FOP marrow cells were immediately injected via retro-orbital injection into syngeneic, same-sex, tamoxifen-induced, busulfan-treated FOP recipient mice to simulate autologous transplantation; control experiments were ones in which FOP marrow cells were electroporated without plasmid delivery (Fig 4A, schematic). After three weeks, marrow reconstitution rates were 92.47% and 95.57% for control and experimental arm with a SD of 3.04% and 1.64% respectively (Fig 4D) as calculated using the illustrated gating strategy. Following 3 weeks of reconstitution, mice received intra-muscular cardiotoxin injection (3 ug x 4 times) into both hindlimbs to stimulate heterotopic bone formation. A second cohort of mice were euthanized 4 weeks after cardiotoxin injection followed by serum ELISA for ActR2A-Fc showing no difference in systemic detection of Fc levels between mice which had received transfected or untransfected FOP bone marrow (Fig 4E, threshold of detection 0.01 ng/mL). Hindlimbs were examined via x-ray (Fig 4F) and microCT (Fig 4G) 4 weeks after local cardiotoxin injury to induce heterotopic bone formation. We found robust heterotopic bone formation in 8 of 10 hindlimbs in FOP mice which received control bone marrow, while we found no bone formation in 11 of 12 hindlimbs with experimental bone marrow (Fig 4H). This relationship was verified based on quantification of heterotopic ossification (HO) on microCT (Fig 4I).

**Figure 4.**
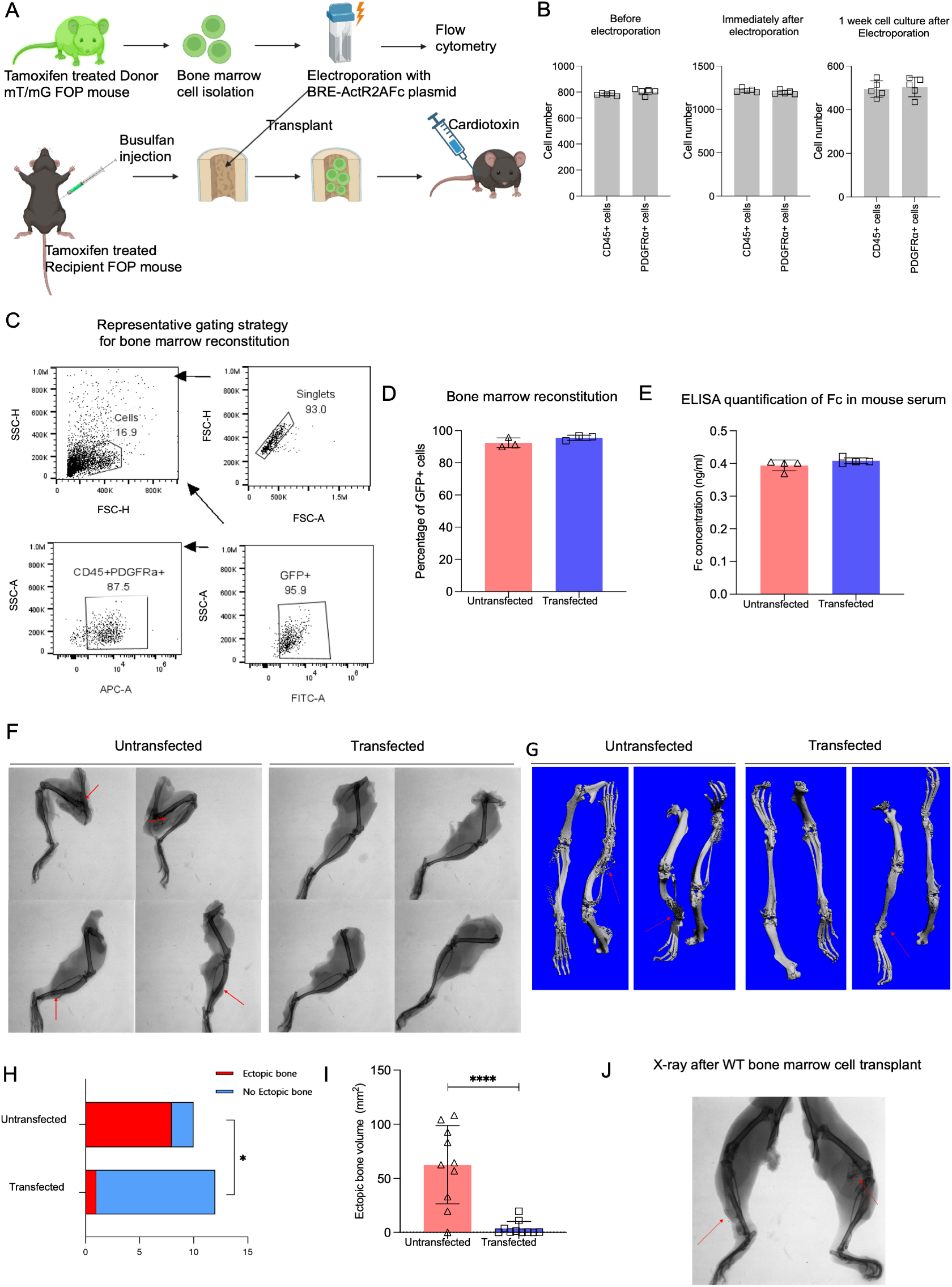
Bone marrow transplantation of transfected cells; 4 weeks post CTX injection. (A) Schematic for in vivo testing of the therapeutic effect of electroporated bone marrow cells. (B) Flow cytometry analysis showing no significant change in the CD45+:PDGFRα+ cell ratio before electroporation, immediately after electroporation or 7 days in cell culture post-electroporation. (C) Flow cytometry gating strategy for donor mouse derived bone marrow cells. (D) The rate of bone marrow reconstitution was high and did not differentiate between mice with transfected vs untransfected bone marrow transplant, showing long-term engraftment of bone marrow cells after 4 weeks of CTX injection. (E) ELISA showed no difference in serum Fc concentration between transfected and untransfected groups indicating absence of systemic ActR2A-Fc production. (F) X-ray images showed higher occurrence of heterotopic bone formation in mice from untransfected group when compared to transfected cohort. (G) Micro-CT reconstruction images showed more heterotopic bone formation in hindlimb of untransfected mice when compared to transfected group. Each dot in the graph represents a single ectopic bone volume in the mouse hindlimb. (H) Chi-squared test showed significant difference in the incidence of heterotopic bone in hindlimbs within untransfected and transfected group, with higher incidence in untransfected group. (I) Quantification of bone volume showed higher volume of heterotopic bone in hindlimb of untransfected mice when compared to transfected group. (J) X-ray images showed heterotopic bone formation in mice transplanted with untransfected wild-type bone marrow cells, confirming that the absence of FOP bone marrow cells alone is insufficient to prevent heterotopic bone formation. N=5 mice for both groups of mice. Each dot in graphs represents a single biological replicate. Statistical significance was established by performing Welsh’s t-test between the means of the groups. * is p<0.05.

To demonstrate that solely correcting the defect in marrow cells would not be sufficient to prevent FOP lesions, we transplanted wild type bone marrow into same-strain FOP recipient mice; mice made FOP bone lesions identifiable on x-ray (Fig 4J). This finding suggests that the therapeutic efficacy observed with our engineered cells is not entirely due to an autocrine inhibition of osteogenic differentiation but includes a paracrine effect of tissue-resident cells. This is consistent with previous findings showing that bone marrow transplantation alone is not sufficient to cure FOP as fibro/adipogenic progenitors (FAPs) are responsible for providing cells for HO^7^.

Next, we sought to verify trafficking of the transplanted marrow to the site of injury, and to verify that the presence of ACVR2A-Fc remained undetectable systemically. Following bone marrow transplantation with GFP+ transfected or untransfected FOP marrow, FOP mice received intramuscular cardiotoxin injection and were euthanized after 1 week; histologic evaluation of hindlimbs demonstrated presence of GFP+ cells, verifying trafficking to the site. However, we noted that GFP+ cells were not present in areas of the hindlimb distant from cardiotoxin injury where there were no inflammatory infiltrates (**Fig 5A**) 27.2% of the total cells at the injury site were positive gor GFP fluorescence. These cells exhibited immunostaining for PDGFRα but not for CD45, indicative of mesenchymal cell fate (**Fig 5B**). The majority of trafficked bone marrow cells also exhibited immunostaining for a HSC panel containing CD11b, TER119, B220, Ly-6G, and CD3 Ab (**Fig 5C**). Tissues from mice treated with transfected and untransfected bone marrow cells were stained for human Fc. Fc immunostain was observed in the muscle tissue at the site of CTX injection but not inside the bone marrow compartment indicating that the secretion of ActR2A-Fc is spatially restricted. Mice that got untransfected bone marrow transplant did not display any signs of Fc stain (**Fig 5D**). After 5 days, average marrow reconstitution rates were 83.97% and 82.4% for control and experimental arm with a SD of 6.38% and 4.41% respectively. (**Fig 5E, F**) The serum of these mice was collected and ELISA performed to measure the concentration of serum ActR2A-Fc. We saw no significant difference in the concentration of serum ActR2A-Fc in mice with transfected and untransfected bone marrow transplant **(Fig 5G**). This shows that the system does not elicit a large non-target response. Most of the activity is restricted to the injured tissue.

**Figure 5.**
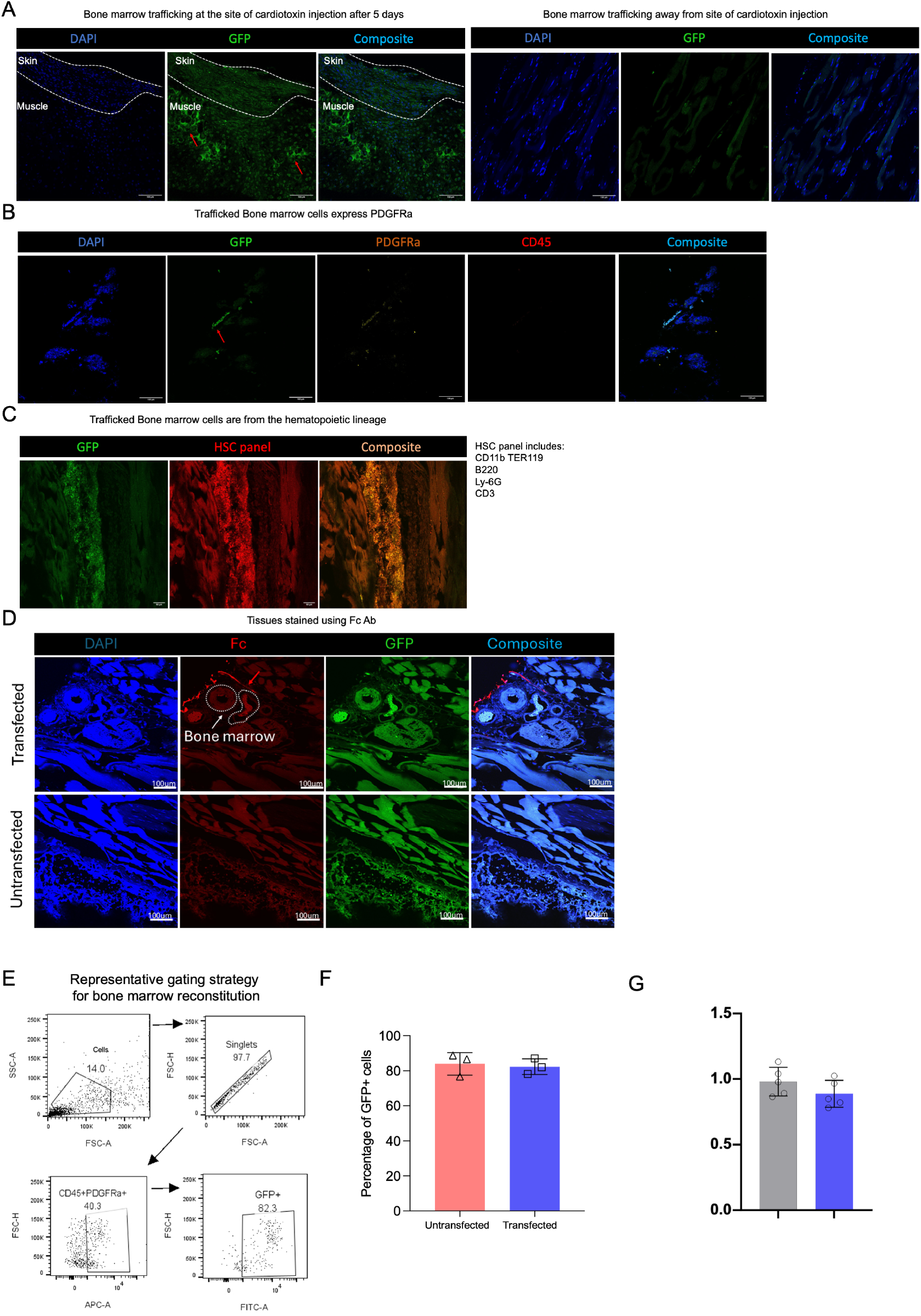
Trafficking of the transplanted bone marrow cells 5 days post CTX injection. (A) Immunofluorescence microscopy showed the presence of GFP+ cells at the site of CTX injection in the recipient indicating trafficking of transplanted donor cells whereas no GFP+ were observed at the site with no CTX injection. 27.2% of the total cells at the injury site were GFP+. (B) Immunofluorescent staining revealed that the trafficked GFP+ cells were also PDGFRα+ cells. (C) Majority of trafficked bone marrow cells also exhibited positive staining for CD11b, TER119, B220, Ly-6G, and CD3 Ab. (D) Fc immunostaining was observed in the muscle tissue but not inside the bone marrow compartment after 5 days of CTX injection in transfected mice whereas no Fc was observed in untransfected mice. (E) Flow cytometry gating strategy for donor mouse derived bone marrow cells. (F) The rate of bone marrow reconstitution was high and did not differ between control mice and treatment mice, showing engraftment as early as 5 days post CTX injection. (G) The serum concentration of ActR2A-Fc did not differ significantly between mice with transfected and untransfected bone marrow transplant. N=5 for each group of mice. Images captured from 3 high powered fields. Each dot in graphs represents a single biological replicate. Statistical significance was established by performing Welsh’s t-test between the means of the groups. * is p<0.05.

### Long term efficacy of engineered bone marrow cells

Next, we sought to assess the efficacy of transplanted engineered bone marrow after 8 weeks of CTX injury. Mice were transplanted with engineered bone marrow and right hindlimb was injected with CTX. Mice were euthanized after 8 weeks and the hindlimbs were scanned using x-rays. Mice that received untransfected bone marrow transplant displayed identifiable FOP bone lesions on x-rays (**Fig 6A**). We noted that the GFP+ transplanted cells were present in the muscle even after 8 weeks of cardiotoxin injection. Most of the GFP+ cells exhibited CD45 staining, indicative of hematopoietic cell fate (**Fig 6B**). Tissue from the site of CTX injection was stained for Fc on the right hindlimb of the mice. Fc immunostain was observed in the right hindlimb at the site of CTX whereas the Fc stain in uninjured left hindlimb was minimal suggesting the local nature of Fc production (**Fig 6C)**. H&E staining of the CTX injection site showed ectopic bone formation in FOP mice with untransfected bone marrow transplant. Transfected mice however showed intact muscle fibers with no signs of ectopic bone. This demonstrated that the trafficking of engineered bone marrow cells can prevent the formation of ectopic bone and provides some early evidence that the structure of the injured tissues may be maintained.

**Figure 6.**
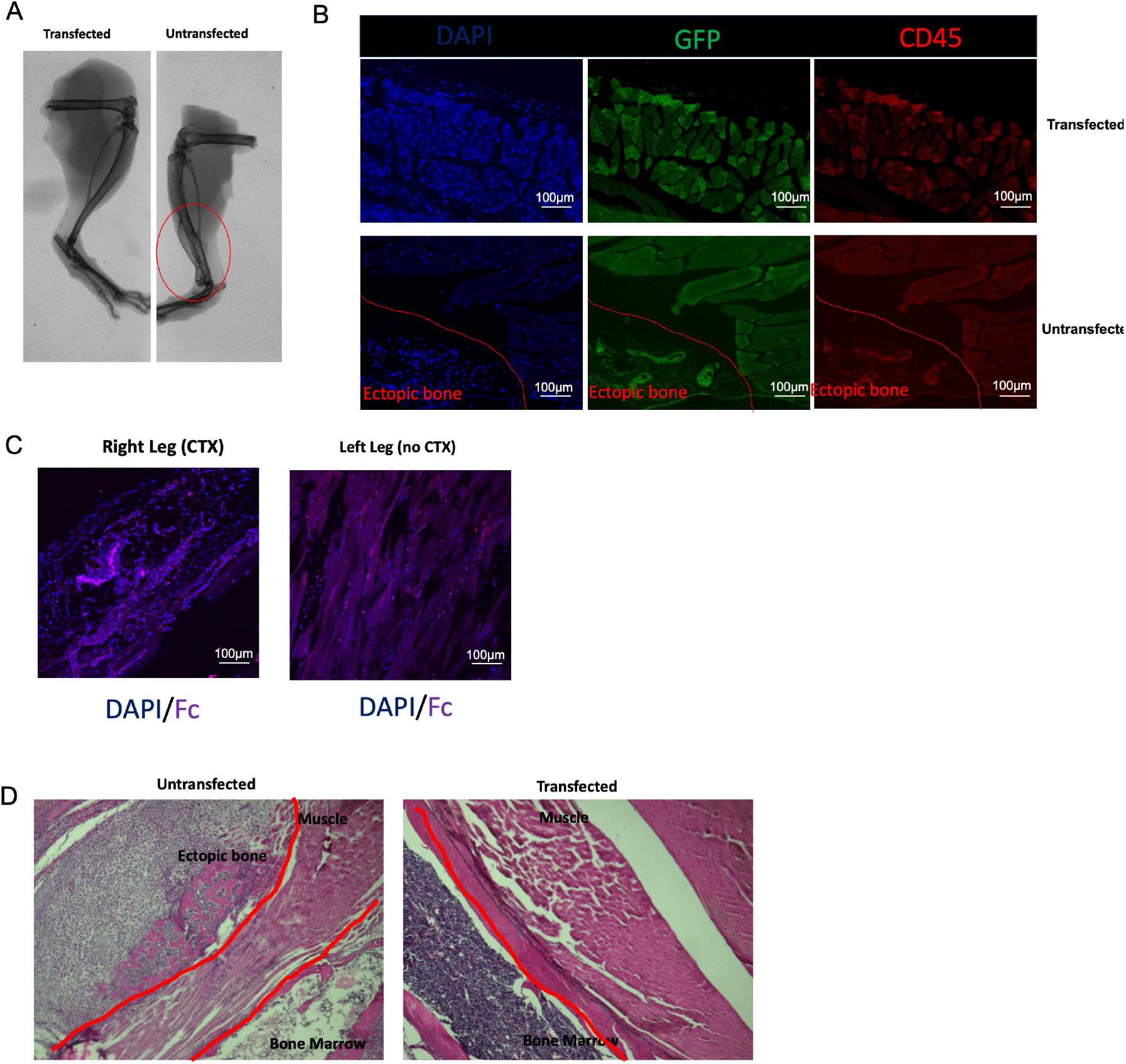
Effect of transplanted bone marrow cells 56 days post CTX injection. (A) X-ray images showed higher occurrence of heterotopic bone formation in mice from untransfected group when compared to transfected cohort after 56 days. All four mice from untransfected group had ectopic bone compared to no ectopic bone in transfected group. (B) Immunofluorescence microscopy showed the presence of GFP+ cells at the site of CTX injection in the recipient indicating trafficking of transplanted donor cells whereas GFP+ cells were observed only in the bone marrow with no CTX injection. The transfected cells were CD45+ cells indicating hematopoietic lineage. (C) Fc immunostaining was observed in the muscle tissue in the right leg after 56 days of CTX injection in transfected mice whereas no Fc was observed in contralateral leg which did not receive CTX injection. (D) H&E staining at the site of CTX injection showed ectopic bone formation in FOP mice with untransfected bone marrow transplant. Transfected mice showed intact muscle fibers with no signs of ectopic bone. N=4 for each group of mice. Images captured from 3 high powered fields.

## Discussion

Here we report on a proof-of-principle closed-loop, autologous cell therapy which is capable of trafficking to sites of injury and activating locally to express an inhibitor of a pathologic ligand. Within the context of FOP, our cell therapy offers a new avenue for drug delivery has the potential to improve site-specific tissue targeting and toxicity profile of the therapeutic agent. Importantly, this technique is amenable only to agents which can be produced by cells in a controlled fashion, such as recombinant peptides. However, our lab has initiated work on cell therapy-based secretion of exosome-packaged therapeutic oligonucleotides as well^13^. In the future, we envision strategies for targeted delivery of chemical agents which may be synthesized within cells, such as steroid hormones or chemotherapeutic agents.

Our work complements other recent advances in cell-based delivery of therapeutic protein against FOP. Prior studies have demonstrated the efficacy of systemic, AAV-mediated delivery to achieve constitutive expression of a ActR2B-Fc in FOP models^14^. Similarly, another approach has utilized engineered mesenchymal cells modified to continuously secrete ActR2B-Fc along with rapamycin, an inhibitor of HO in FOP^15^. While these strategies are promising, they rely on the continuous and systemic production of a biologic agent. Our closed-loop approach is predicated on the ability to detect the presence of a ligand associated with the disease and produce the therapeutic protein in response locally and without a detectable systemic presence. We surmise that by being responsive to local levels of the offending ligand, the cell therapy is finely tuned to the microenvironment, preventing large amounts of gene expression which would result in systemic “spillage”. Indeed we did not identify systemically elevated levels of ActR2A-Fc and we did not identify any obvious systemic effects associated with intravenous infusion of the drug such as hemorrhage or telangiectasias. However, further studies are required to rigorously determine whether these may develop over longer periods of time. We are encouraged by the absence of systemically quantifiable ActR2A-Fc levels however.

The therapeutic system we have developed is distinct in that it is designed as a closed-loop circuit. In our model, the expression of ActR2A-Fc is not constitutive; rather, it is induced specifically by the local, pathogenic BMP signaling that characterizes an inflammatory flare leading to heterotopic ossification (HO). Previously, BRE was shown to be uniquely activated by Activin A in cells harboring the ActR1 *R206H* mutation, but not in wild type cells^8^. Our validating experiments for the BRE promoter selection indicated that upon a decrease in Activin A levels, there is a concomitant decrease in expressivity. This lends itself to closed-loop expression which is important given genomic permanence of the transposon. As our results demonstrate, the engineered marrow cells traffic to the site of injury, ensuring that the therapeutic is produced precisely where and when it is needed. This spatiotemporal control is a key differentiator from constitutive delivery systems and may offer a significant advantage by minimizing systemic exposure, thereby potentially enhancing the long-term safety profile of the treatment.

Our selection of ActR2A-Fc as the therapeutic peptide of interest is based on our knowledge of its Activin A binding capacity. Other candidates including ActR2B-Fc may be considered in the future as well. While structurally different from these peptides, anti-Activin A antibodies (*e*.*g*. Garetosmab) may be considered in the future. In this way, our cell therapy may behave similar to a pre-programmed B-cell therapy which is capable of resident in the marrow. In this manuscript, we did not compare ActR2A-Fc with Garetosmab with respect to efficacy or adverse effects; we believe that peptide selection is important, but in this proof-of-concept study is secondary to the development and deployment of a guiding therapeutic framework.

Our cell therapy also provides a practical benefit over exogenous peptide delivery and oral therapies like orally administered ALK2 kinase inhibitors^16^. By enabling endogenous production of the recombinant peptide, unique peptide manufacturing, storage/shipping, and injection requirements are obviated. Instead of ex vivo bioreactor-based manufacturing with complex purification requirements, the burden of manufacturing is instead directed towards standard DNA synthesis. Systemic infusion of the agent is no longer required due to the ability of these cells to traffic to the desired site. Technical constraints for local injection are also bypassed as cells directly migrate to the affected site, and we can take advantage of the power of a cell therapy system capable of detecting and responding to the earliest indicators of disease even before typical signs/symptoms are observable by the patient or provider. Autologous BMT is a significant undertaking. Ultimately, the viability of a BMT-based approach for FOP will depend on a clear demonstration of profound, lifelong efficacy and a safety profile that outweighs both the acute risks of the transplant procedure and the convenience of chronically administered alternatives.

Our strategy offers numerous opportunities for iterative improvements and modifications. For FOP, one may consider modifying promoter selection from BRE to another promoter which may have improved expressivity or specificity for the site of interest. Importantly, selection of promoter is a driver for not only specificity but intensity of drug expression. This presents an immense opportunity to leverage the trove of transcriptional data which exists for myriad diseases to then identify optimal disease-specific promoters. The nature of disease initiation may be re-defined to the activation of a specific promoter which is sufficient to indicate the appropriate temporal window during which to treat the patient. One may also alter the desired product from ActR2A-Fc to another peptide of interest opening the door to myriad different peptide therapeutics being considered without the costly infrastructure required for peptide synthesis and purification. Most importantly, this strategy is amenable to use across different diseases for which systemic and long-term drug delivery are not acceptable due to adverse effects. As we continue to develop this technology, we will address these permutations and examine them for therapeutic viability using in vivo models.

## STAR METHODS

### KEY RESOURCES TABLE

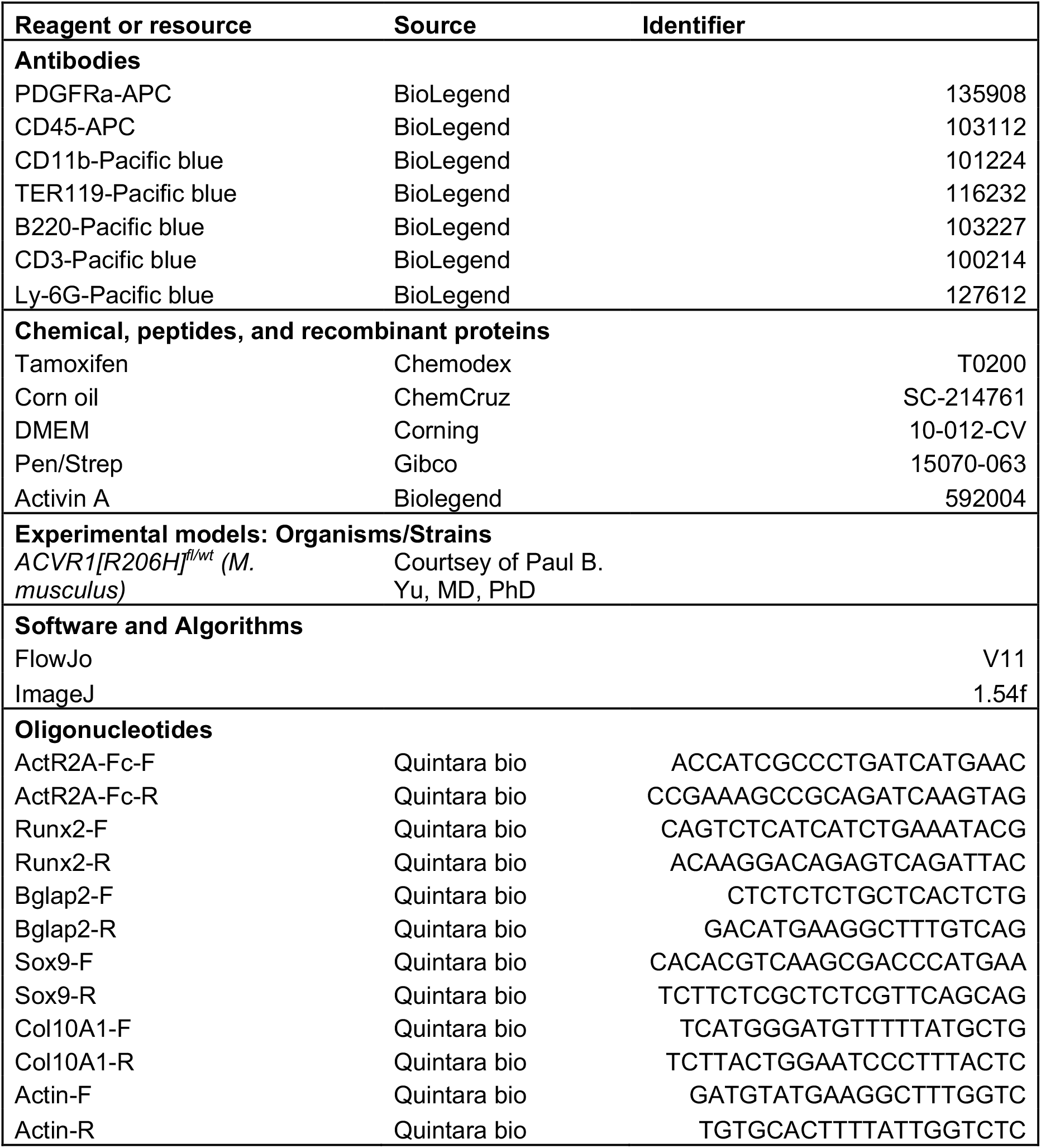

### EXPERIMENTAL MODEL AND STUDY PARTICIPANT DETAILS

#### Mice

The experiments and handling of mice were performed in accordance with the protocol approved by Institutional Animal Care and Use Committee (IACUC) of Brigham and Women’s Hospital (BWH, Protocol number 2020N000036).

*ACVR1[R206H]*^*fl/wt*^ mice^8^ were mated with Ubc.CreERT2;*Rosa26*^*mTmG*^ mice^17^ to generate Ubc.CreERT2;*Rosa26*^*mTmG*^*;ACVR1[R206H]*^*fl/wt*^ and Ubc.CreERT2; *ACVR1[R206H]*^*fl/wt*^ mice. Ubc.CreERT2;*Rosa26*^*mTmG*^*;ACVR1[R206H]*^*fl/wt*^ mice severed as donors for bone marrow transplant whereas Ubc.CreERT2; *ACVR1[R206H]*^*fl/wt*^ mice were recipients. These were maintained in heterozygosity on a C57BL/6 background. Both male and female mice were used between 6 and 12 weeks of age. All mice were age and sex matched between groups.

### METHOD DETAILS

#### Tamoxifen treatment

Tamoxifen was intraperitoneally injected into UBC-Cre-ERT2;*ACVR1*[*R206H*] mice at the age of 6 weeks twice at a interval of 3 days. Tamoxifen (T0200, Chemdex) was dissolved in corn oil (SC-214761, Chemcruz) at a concentration of 20mg/ml, 100 μl was injected.

#### Activin A Reporter Vector Construction

The BMP Recognition Element (BRE) was used upstream of the minimal CMV promoter and ligated with the ActR2A-Fc encoding gene. The product was inserted into a PiggyBac^18^ vector containing Blasticidin-resistance gene for selection (VectorBuilder). E. coli DH5α cells carrying the plasmid was cultured in LB medium containing 100μg/ml ampicillin, and the plasmid was isolated using ZymoPURE II plasmid maxiprep kit (D4202, Zymo Research).

#### Cell culture

HEK293 cells were cultured in DMEM (10-013-CV, Corning) supplemented with 10% fetal bovine serum (Sigma) and 1% penicillin/streptomycin (15070-063, Gibco). Bone marrow cells were isolated from mice. Specifically, tamoxifen treated donor FOP mice were euthanized using cervical dislocation and both hindlimbs were obtained with femur, tibia, and fibula intact. Muscle and tissue residues around the femur were removed, and the proximal end of the femurs were cut using sharp scissors. A 10 cc syringe with a 21-gague needle was filled with ice-cold PBS and inserted into the distal end of the femur. PBS was injected into the femur to flush the bone marrow out from the proximal end onto a 70 μm nylon cell strainer. The cells were collected in a 50 ml Falcon tube and centrifuged at 1500 rpm for 5 minutes at 4 °C. The cells were resuspended in ice-cold PBS and electroporated immediately. After electroporation, cells were cultured in a growth medium containing DMEM, 10% FBS and 1% penicillin/streptomycin. All cells were cultured in a humidified atmosphere with 5% CO2 at 37 °C.

#### Transfection

HEK293 cells were transfected using Lipofectamine 3000 (ThermoFisher). Transfections were performed in 6-well plates (1×10^6^ cells/well). For each well, 4 μg vector was mixed with 1 μg pBASE plasmid carrying PiggyBac transposase. The transfection mix was prepared according to the manufacturer’s protocol and added into the cell culture media. Cells were washed with fresh culture media after 48 hours and 100ng/ml Activin A (592004, Biolegend) was added to half of the wells. Cells were harvested after 1 week and the ability to produce ActR2A-Fc mRNA and protein was examined by RT-PCR and ELISA.

#### Electroporation

Bone marrow cells were counted using a hemacytometer. Appropriate dilution was performed in PBS to achieve a concentration of 1×10^4^ cells/μL. 500 μL cell suspension was added to a 2 cm gap electroporation cuvette along with 15 μg BRE-ActR2A-Fc vector and 5 μg pBASE transposase plasmid and chilled on ice for 15 minutes. A single 5 ms pulse of 200 volts was used to electroporate (GenePulser Xcell, BioRad) the cells and the cuvette was placed on ice for another 15 minutes. 1.5 ml DMEM/FBS/Pen/Strep was added to the cells and plated in a 6-well plate. Cells were washed with fresh culture media after 48 hours and 100 ng/ml Activin A was added to half of the wells. Cells were harvested after 1 week and the ability to produce ActR2A-Fc mRNA and protein was examined by RT-PCR and ELISA.

#### RT-qPCR

Total mRNA was isolated from the transfected cells using Direct-zol RNA Miniprep Kits (Zymo Research). Reverse transcription was performed using qMax cDNA Synthesis Kit (Accuris) with 100ng of starting RNA. Real-Time PCR was performed using the following primers for *ActR2A-Fc, Runx2, Bglap2, Sox9, Col10A1*, and *Actin* respectively. The PCR condition was: 95 °C for 10 minutes; 40 cycles 95°C for 15 s, 60 °C for 1 minute, 72 °C for 30 s; 72 °C for 10 min The PCR was performed using the machine QuantStudio 7 Flex Real-Time PCR Systemand analyzed using QuantStudio™ Real-Time PCR Software. B-Actin was used as the house keeping gene and gene expression was measured using the ΔΔCt method.

#### ELISA

Blood was collected from mice using cardiac puncture, followed by incubation for 30 minutes at room temperature. The clot was removed by centrifuging the tube at 1000g for 10 minutes and the supernatant was collected. The supernatant was stored at -80°C until further use. Human Fc ELISA kit (EK000095-HUFC-2, Syd labs) was used to measure Fc concentration in the serum. Similarly, culture media was collected after 7 days of cell culture. Human Fc ELISA kit was used to measure Fc concentration in the culture medium.

#### In vivo mouse model of FOP

The model was initiated by replacing wild-type *ACVR1* sequence with its *R206H* counterpart into the sense strand. This was achieved by treating the UBC-Cre-ERT2;FOP mice with tamoxifen (100mg/kg, intraperitoneally) in corn oil for 4 days, which activated CreERT2. The same injection also replaced tdTomato-encoding exon with EGFP-encoding exon into the sense strand in UBC-Cre-ERT2;FOP;mTmG mice. To ablate bone marrow cells from the recipient mice, 25mg/ml stock of busulfan in DMSO was diluted in 0.2% saline and injected into the mice (20mg/kg, intraperitoneally) for 4 days. For bone marrow transplant, donor bone marrow cells were isolated from tamoxifen-treated UBC-Cre-ERT2;FOP;mTmG mice. Bone marrow was flushed with PBS, filtered through a 70 μm cell strainer, and electroporated with or without the BRE-ActR2A-Fc vector. A total of 1 × 10^6^ cells were then injected retro-orbitally into recipient UBC-Cre-ERT2;FOP mice. Mice were anesthetized using isofluorane before the injection. Three weeks after the bone marrow transplant, cardiotoxin (0.1 mg/ml in PBS) was injected intramuscularly into the hindlimb of the mice to induce muscle injury and promote heterotopic ossification.

#### Micro-CT and anatomical measurements

Left and right hind limbs were collected using high-resolution Micro-Computed Tomography (μCT40, SCANCO Medical AG, Brüttisellen, Switzerland). Scan parameters were 30 μm^3^ isotropic voxel size, 55 kVp peak X-ray tube intensity, 145 μA X-ray tube current, and 300 ms integration time. Resultant DICOM images were exported for measurement of heterotopic bone volume.

#### Flow Cytometry

Muscle and tissue residue around the femur were removed, and the proximal end of the femurs was cut using sharp scissors. A 10 cc syringe with a 21-gauge needle was filled with ice-cold PBS and inserted into the distal end of the femur. PBS was injected into the femur to flush the bone marrow out from the proximal end onto a 70 μm nylon cell strainer. The cells were collected in a 50 ml Falcon tube and centrifuged at 1500 rpm for 5 minutes at 4°C. The cells were then resuspended in ice-cold PBS and counted using a hemacytometer. Appropriate dilution was performed to achieve a concentration of 1×10^4^ cells/μL. The cells were resuspended in blocking buffer (5% goat serum) for 1 hour. The cells were washed twice in PBS and the pellet was suspended in APC-PDGFRa (135908, Biolegend) or APC-CD45 (103112, Biolegend) for 1 hour. The cells were washed in PBS to remove the unconjugated antibodies and analyzed using Beckmac Coulter with appropriate filters. The data was analyzed using FlowJo. Live single cells were gated based on FSC/SSC followed by FSC-A/FSC-H for singlet gating. Singlets were gated with marker-specific gates.

#### Immunofluorescence staining

Tissue was fixed in paraformaldehyde for 48 hours and washed with PBS. The fixed tissue was frozen in OCT using liquid nitrogen and stored in -80°C before cutting. The blocks were cut using a cryostat with the tissue thickness of 10μm and mounted on a positively charged slides. Tissue sections were blocked with 10% goat serum, 1% bovine serum albumin (BSA), and 0.3% Triton (MilliporeSigma) in PBS for 1 hour and incubated with primary antibodies at 4°C overnight. The sections were probed with antibodies for APC-PDGFRα (1:100). The HSC panel included CD11b, TER119, B220, CD3, Ly-6G (101224,116232, 103227, 100214, 127612, Biolegend) conjugated with pacific blue. Sections were washed for 3 x 5 min in PBS. ProLong Diamond Antifade Mountant with DAPI (P36971, Invitrogen) was used to stain the nuclei and mount the samples. Fluorescent images were taken by Olympus FluoView confocal microscope at original magnification, 20×. Quantification of immunofluorescence staining was performed by using ImageJ software.

### QUANTIFICATION AND STATISTICAL ANALYSIS

Data are presented as mean ± S.D. Statistical analysis were performed using GraphPad Prism 9. Comparisons between two groups were performed by using 2-tailed Student’s t-tests. Comparisons among multiple groups were performed by using a one-way ANOVA with a Tukey’s multiple-comparisons test. Significant difference was established when P values was smaller than 0.05.

## Acknowledgments

This work is supported by NIH/NIAMS grant 5R21AR077769 (SA), NIH/NIAMS grant 1K08AR082031 (SA), Plastic surgery foundation National endowment for plastic surgery award (SA).

## Notes

### Competing Interest Statement

The authors have declared no competing interest.

